# Selective B cell depletion upon intravenous infusion of replication-incompetent anti-CD19 CAR lentivirus

**DOI:** 10.1101/2020.05.15.098335

**Authors:** Craig M. Rive, Eric Yung, Lisa Dreolini, Scott D. Brown, Christopher G. May, Daniel J. Woodsworth, Robert A. Holt

**Affiliations:** Canada’s Michael Smith Genome Sciences Centre, BC Cancer, Vancouver, BC, V5Z 1L3, Canada; Department of Medical Genetics, University of British Columbia, Vancouver, BC, V6T 1Z4, Canada; Molecular Biology & Biochemistry, Simon Fraser University, Burnaby, BC, V5A 1S6, Canada

**Keywords:** CAR-T, Immune effector cells, viral vector, gene transfer, gene therapy, leukemia, lymphoma

## Abstract

Anti-CD19 CAR-T therapy for B cell malignancies has shown clinical success, but a major limitation is the logistical complexity and high cost of manufacturing autologous cell products. If engineered for improved safety, direct infusion of viral gene transfer vectors to initiate *in vivo* CAR-T transduction, expansion and anti-tumor activity could provide an alternative, universal approach. To explore this approach we administered approximately 20 million replication-incompetent VSV-G lentiviral particles carrying an anti-CD19CAR-2A-GFP transgene comprising either an FMC63 (human) or 1D3 (murine) anti-CD19 binding domain, or a GFP-only control transgene, to wild-type C57BL/6 mice by tail vein infusion. The dynamics of immune cell subsets isolated from peripheral blood were monitored at weekly intervals. We saw emergence of a persistent CAR-transduced CD3^+^ T cell population beginning week 3-4 that reached a maximum of 13.5 +/-0.58% (mean +/-SD) and 7.8 +/-0.76% of the peripheral blood CD3^+^ T cell population in mice infused with ID3-CAR or FMC63-CAR lentivector, respectively, followed by a rapid decline in each case of the B cell content of peripheral blood. Complete B cell aplasia was apparent by week 5 and was sustained until the end of the protocol (week 8). No significant CAR positive populations were observed within other immune cell subsets or other tissues. These results indicate that direct IV infusion of conventional VSV-G pseudotyped lentiviral particles carrying a CD19 CAR transgene can transduce T cells that then fully ablate endogenous B cells in wild type mice.

## Introduction

Ordinarily, a cancer patient’s immune system can recognize cancer cells as being altered, and mount a natural immune response. T cells are known to be the main mediators of the anti-cancer immune response but sometimes the natural T cell response is ineffective; T cells may fail to recognize tumor cells, fail to activate, or fail to sustain a response long enough to have an impact. It is possible for these limitations to be overcome by genetic engineering strategies that involve isolating a sample of a patient’s T lymphocytes, genetically modifying and activating the cells *ex vivo*, and then re-administering them to the same patient. The genetic modification step involves introducing into the T cells an extra gene that carries instructions for a new antigen receptor that may be, for example, a recombinant alpha-beta T cell receptor (αβTCR) or a Chimeric Antigen Receptor (CAR). Much of the growing interest in genetically engineered IECs as a new class of therapeutic is driven by the impressive results CAR-T therapies have shown in patients with relapsed/refractory B-cell Acute Lymphoblastic Leukemia or Non-Hodgkin Lymphoma. These patients, for whom standard therapies have failed, have historically had dismal outcomes with survival measured in weeks or months. Remarkably, CAR-T therapy targeting CD19 has led to meaningful and durable remissions for large proportion of these patients in many different studies[1–7].

Currently, most genetically engineered Immune Effector Cell (IEC) therapies require several steps including i) manufacturing plasmid encoding the synthetic transgene and ancillary plasmids, ii) delivery of these plasmids into producer cells to generate virus, iii) collection of patient autologous T cells and integration of the transgene into the genomes of these cells by viral transduction, iv) expansion of the transduced T cells *ex vivo* to produce the therapeutic cells, v) extensive release-testing of the manufactured product, vi) preconditioning lymphodepletion of the patient, vii) infusion of the modified autologous cells, and viii) intensive clinical and immunological monitoring. Thus, there is considerable infrastructure and expertise required to deliver IEC treatments safely and successfully. The complexity of manufacturing IECs is a critical limitation to the cell therapy field, and a major contributor to the high cost of new cell-based interventions coming to market. Hence, there is considerable interest in the potential for “universal” alternative to autologous CAR-T products, with some encouraging results seen from gene-edited allo-CAR-Ts pre-clinically [8] and in case studies[9,10]. Likewise, the immortalized IEC cell line NK-92 has shown encouraging safety and efficacy in a recent phase 1 trial, but the requirement of irradiating the cells pre-infusion to reduce oncogenic risk will likely remain limiting for this approach.[11] The generation of a bank of allogeneic CAR-T cells by *in vitro* differentiation and maturation of hematopoietic stem and progenitor cell precursors is an alternative and highly compelling strategy. Demonstrated in principle, this approach awaits further optimization.[12]

In principle, universal IEC therapy could also be achieved by direct infusion of a gene transfer vector, such that cell manufacturing is sidestepped completely. Different approaches to *in vivo* gene delivery have been explored, preclinically, for CAR-T therapy. Nanoparticles that carry a CAR transposon and that are coated with anti-CD3 targeting antibodies can transduce T cells *in vivo* yielding functional CAR-T cells[13], as can lipid nanoparticles that carry CAR mRNA and display anti-CD5 targeting antibodies[14]. Similarly, *in vivo* delivery of high doses of 10^10^ to 10^11^ lentiviral particles that bear CAR transgenes, and that have been pseudotyped to target either CD3+, CD8+ or CD4+ T cells, can yield functional CAR-T[15–17]. Here, we show efficient *in vivo* generation of functional CAR-T cells in wild-type mice upon infusion of lower doses (on the order of 10^6^ to 10^7^ particles) of conventional VSV-G lentivirus bearing transgenes that encode conventional FMC63 or ID3 anti-CD19 binding domains.

## Results

During our early pre-clinical development of an anti-CD19 CAR intervention for B cell malignancies we explored how CAR-T cells behave *in vitro* in the context of antigen driven expansion. Using a standard 2^nd^ generation anti-CD19 CAR design[18] (EF1α promoter, FMC63 scFv CD19 binding domain, CD8α hinge and transmembrane domain, 4-1BB co-stimulatory domain, CD3z signaling domain and 2A-GFP tag), we observed, as have many other investigators, that B cells drive robust expansion of functional CD19 CAR-T cells *in vitro*. This led us to explore *in vivo* transduction and expansion of CAR-T cells *in vivo* using a murine model system. We constructed a CAR where we changed the scFv binding domain from FMC63[19] (a monoclonal antibody raised against human CD19) to 1D3[20] (a monoclonal antibody raised against mouse CD19). To verify activity *in vitro*, 2.5×10^5^ splenocytes from wildtype C57BL/6 mice were transduced with lentivirus encoding either the 1D3-CD19CAR-GFP (murine), FMC63-CD19CAR-GFP (human) or GFP-only transgene, then stimulated with a further excess of irradiated autologous murine splenocytes as a source of murine CD19 antigen. We observed that murine T-cells modified with either of the CARs expanded rapidly and selectively. Specifically, for murine T cells modified with the human CAR, expansion was 8.8+/-0.03 (mean +/-SD) fold at day 63 relative to GFP transduced T cells (p < 0.004 paired, two-tail T test), and for murine T cells modified with the murine CAR expansion was 12.9 +/-0.09 fold at day 63, relative to GFP transduced T cells (p < 0.002 paired, two-tail T test). The murine control T cells transduced with GFP-only lentivirus showed very limited yet detectable expansion, possibly driven by the original CD3/CD28 activation prior to transduction [**Figures 1A and 1B**]. All expanded murine CAR-T cells showed robust and selective killing of murine B cells *in vitro*. Target murine B cells co-cultured with either effector 1D3-CD19CAR-GFP or FMC63-CD19CAR-GFP modified murine T cells showed 24.8 +/-1.69% (mean +/-SD) and 18.5 +/-1.08% of murine B cells were unviable (FVS780 positive) after the 5 hour co-culture, respectively [**Figure 1C**].

**Figure 1:**
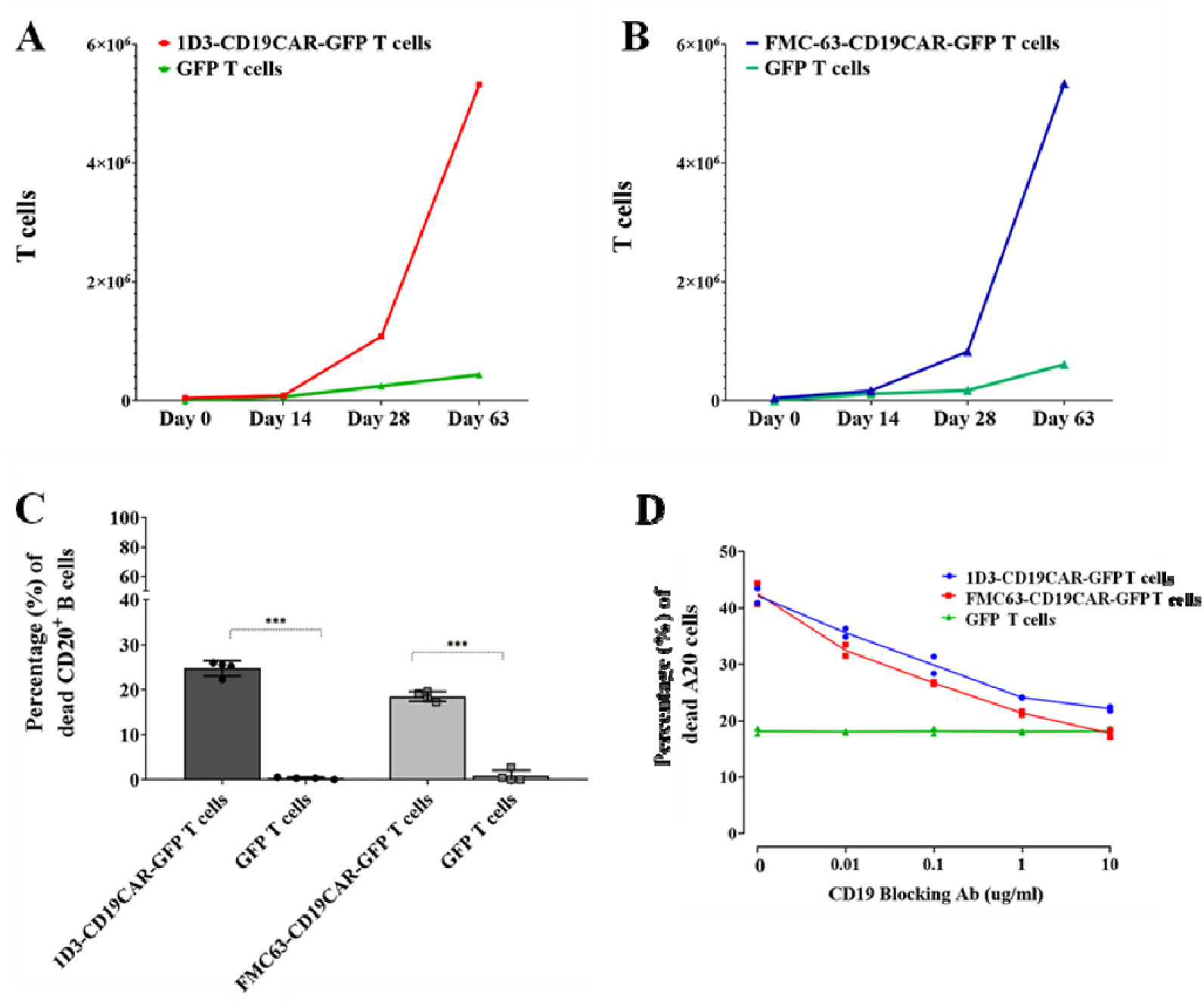
*In vitro* proliferation and cytolytic activity of murine CAR-T cells. Figure 1A. 2.5×10^5^ C57BL/6 splenocytes were transduced with 1D3-CD19CAR-GFP or GFP-only lentivirus at an MOI of 5. The transduced splenocytes were then co-cultured with an excess of irradiated C57BL/6 splenocytes which provided a source of B cells expressing the CD19 antigen. The 1D3-CD19CAR transduced splenocytes showed a 12.2+/-0.09 (mean +/-SD) fold expansion (red, square), compared to GFP transduced (green, triangle) (p < 0.002 paired, two-tail T test). **Figure 1B**: 2.5×10^5^ C57BL/6 splenocytes were transduced with the FMC63-CD19CAR-GFP or GFP-only lentivirus at an MOI of 5. Splenocytes were then co-cultured with irradiated C57BL/6 splenocytes which provided a source of B cells expressing the CD19 antigen. The FMC63-CD19CAR transduced splenocytes showed a 8.8 +/-0.03 fold expansion (orange, square), compared to GFP transduced (cyan, triangle), (p < 0.004 paired, two-tail T test). **Figure 1C**: 1×10^5^ FMC63-CD19CAR-GFP or 1D3-CD19CAR murine T cells from day 63 of the expansion assays were co-cultured for 5 hours with target B cells, isolated from C57BL/6 splenocytes, at a E:T ratio of 1:1. Unviable target murine B cells were measured through the shift in the FVS780 signal, measured by flow cytometry [**Figure S4**]. Expanded murine 1D3-CAR-T cells showed robust and selective targeting of murine B cells, with 24.8 +/-1.69% of B cells being unviable after a 5 hour co-culture. The FMC63-CD19CAR T cells also showed robust and selective targeting of murine B cells with 18.5 +/-1.08 % of B cells being unviable after a 5 hour co-culture.. Two-way ANOVA, Sidak’s multiple comparison ***p<0.002. **Figure 1D**: 1D3-, FMC63-CD19CAR, and GFP-transduced and expanded murine T cells were co-cultured with target A20 cells for 4 hours. The CD19 antigen on the A20 cells, were blocked with either the CD19 -1D3 or -FMC63 blocking antibody at a starting concentration of 10μg/ml decreasing 10 fold down to 0μg/ml, at an E:T ratio of 4:1. Using flow cytometry, target cell death was measured through the shift in the FVS780 signal, [**Figure S3** and **Table S1**]. The murine CAR-T cells showed robust and selective response when co-cultured with A20 cells. This response decreased with increasing CD19 blocking antibody concentration. 38.8+/-0.71% of unblocked A20 cells were unviable when co-cultured with the 1D3-CD19CAR T cells, which decreased to 19.7 +/-0.28% of A20 cells blocked with 10μg/ml of 1D3-CD19 blocking antibody. 41.1+/-0.92% of unblocked A20 cells were unviable when co-cultured with the FMC63-CD19CAR T cells, which decreased to 20.8 +/-0.57% of A20 cells blocked with 10μg/ml of FMC63-CD19 blocking. Two-way ANOVA p = 0.04. Nonlinear regression analysis (Sigmoidal) analysis of 1D3-CD19CAR T cells vs A20 CD19 blocking antibody dose response, R^2^ =0.95. FMC63-CD19CAR T cells vs A20 CD19 blocking antibody dose response, R^2^ = 0.96. GFP vs A20 CD19 blocking antibody dose response, R^2^ =0.01.

For further validation we used 1D3 and FMC63 CD19-blocking antibodies to assess murine CAR-T cell specificity for the mouse CD19 antigen expressed by the A20 cell line [**Figure 1D**], As a baseline, 38.8 +/-0.71% (mean +/-SD) of the target A20 cells were unviable after a 4 hour co-culture with the effector 1D3-CD19CAR-GFP modified murine T cells with no blocking antibody present. Upon titration of 1D3 blocking antibody, the ability of the 1D3-CD19CAR-GFP modified murine T cells to target the A20 cells produced a decrease in unviable A20 cells to 19.7 +/-0.28% (mean +/-SD), which was the level of background killing observed for murine T cells transduced with the GFP-only transgene, likely due to alloreactivity. Likewise, 41.1 +/-0.92% (mean +/-SD) of the target A20 cells were unviable after a 4 hour co-culture of the effector FMC63-CD19CAR-GFP modified murine T cells with no blocking antibody present and 20.8 +/-0.57% of the A20 cells upon titration of FMC63 antibody. Interestingly, these results confirm that the FMC63 scFv is able to bind murine CD19 to the degree necessary for functional CAR-T activation in this experimental system. To our knowledge this has not previously been reported.

We then administered lentivirus encoding each CAR or GFP-only lentivirus to C57BL/6 wild type mice (age 6-8 weeks) by tail vein infusion, and drew peripheral blood at regular intervals. For the following experiments, the number of mice per treatment group was 8 (n=8). In **Figure 2** and **Figure 4**, error bars represent the standard deviation away from the calculated mean of each treatment group. We observed that GFP-expressing T cells appear in the blood of mice 3 weeks after the initial injection of 3.6-4×10^6^ infectious units (IU) of either the 1D3-or FMC63-CD19CAR-GFP lentivirus, but not after injection of the same dose of GFP-only lentivirus **[Figure 2]**. Specifically, we saw emergence of a persistent 1D3-CD19CAR-GFP transduced CD3^+^ T cell population beginning at week 3 that reaching a maximum of 13.5 +/-0.58% (mean +/-SD) of the peripheral blood CD3^+^ T cell population by week 5 [**Figure 2A-2C**]. This coincided with the loss of B cells, seen 5 weeks after the initial injection **[Figure 2C]**. In experiments where we infused murine T cells modified with the FMC63-CD19CAR-GFP construct we also saw the emergence of a persistent FMC63-CD19CAR-GFP transduced CD3^+^ T cell population beginning at week 5 that reached a maximum of 7.8 +/-0.76% of the peripheral blood CD3^+^ T cell population by week 5 **[Figure 2A-2C]**. Again, this coincided with a loss of B cells, seen 5 weeks after the initial injection. This further confirms functional recognition of murine CD19 by FMC63 **[Figure 2C]**. With either CAR, B cell aplasia was sustained until the end of the protocol (week 8). Mice treated with 3.6-4×10^6^ IU of the GFP control lentivirus had no GFP expressing T cells and did not show the same B cell loss **[Figures 2A-2C]**. We looked for GFP expression in immune cell subsets other than T cells, including the CD20/CD45R/B220 B cell population, CD11b^+^ macrophage, and CD335^+^ natural killer (NK) cells, but found that only T cells expressed the reporter GFP gene, indicative of CAR expression **[Figure 2D,2E]**. Our interpretation of these results is that interaction between the *in vivo* transduced CAR-T cells and endogenous B cells drives CAR-T cell expansion and CAR-T mediated B cell depletion, resulting eventually in B cell aplasia. The only overt change to mice treated with either 1D3-or FMC63-CD19CAR lentivirus was a modest decrease in weight of 5.5 +/-2.97% (mean +/-SD) and 3.6 +/-1.78%, respectively over the first week after treatment [**Figure S6**].

**Figure 2:**
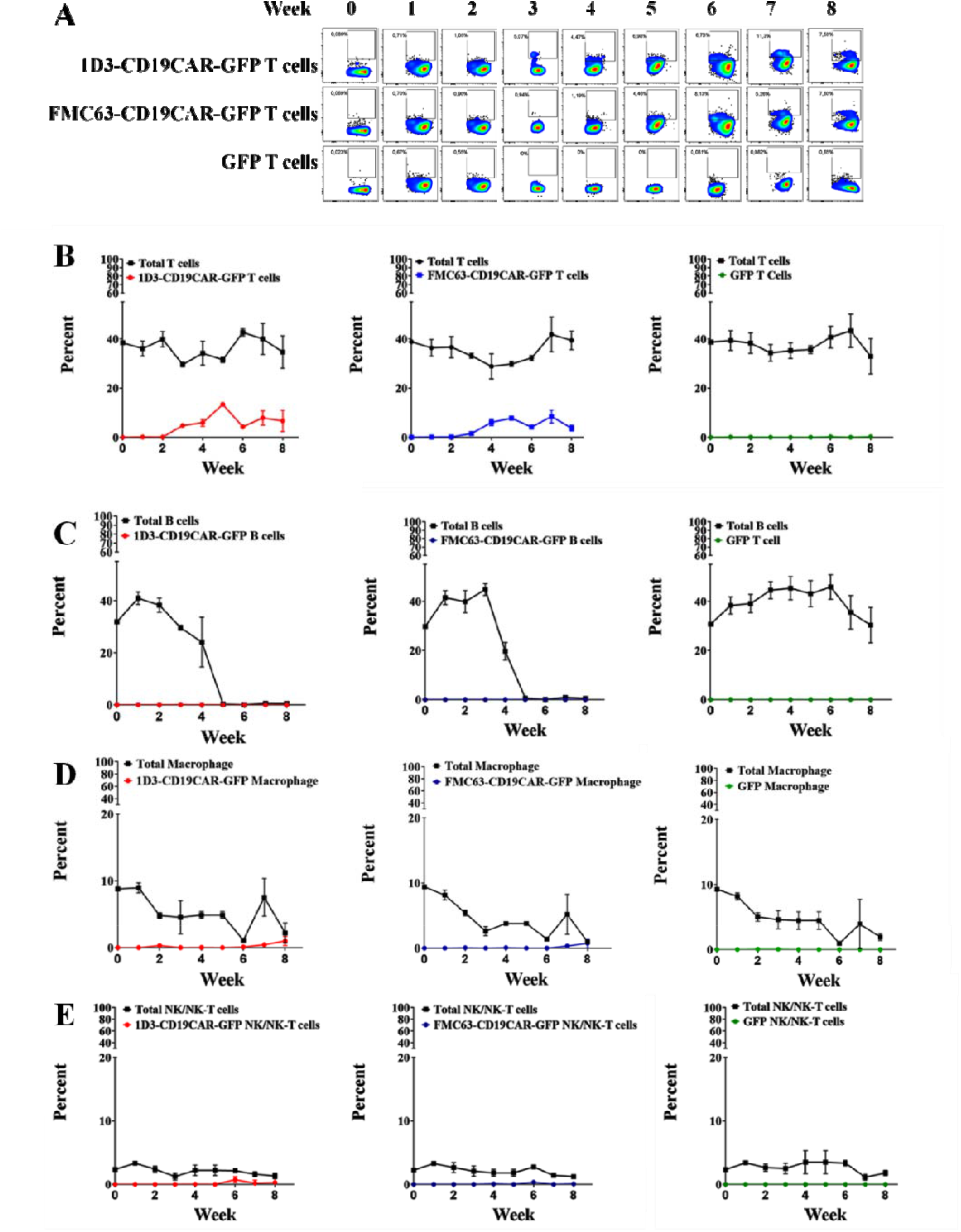
The abundance of immune cell subsets in treated mice over time, as a percent of total PBMC. C57BL/6 wildtype mice were treated with a single IV injection of lentivirus (3.6-4.0 x10^6^ IU in 200μl PBS), to deliver either the 1D3-CD19CAR-GFP CAR, FMC63-CD19 CAR, or control (GFP-only) N equals 8 mice per group and error bars represent the standard deviation from the mean value reported for each group. T cells, but not other immune cell subsets tested, show a substantial transduced cell population. **Figure 2A**: Representative flowgrams showing increasing GFP positive T cells in PBMC of treated mice, over time. The X-axis of the flowgrams is CD3 positivity and GFP positivity on the Y-axis. from **Figure 2B-2E. Figure 2B**: Total T cells versus 1D3-, FMC63-CD19CAR, and GFP transduced T cells (CD3^+^, CD90.2^+^ T cells). **Figure 2C**: Total B cells versus 1D3-, FMC63-CD19CAR, and GFP transduced B cells (CD20^+^, CD45R/B220^+^ B cells). **Figure 2D:** Total macrophage versus 1D3-, FMC63-CD19CAR, and GFP transduced macrophage (CD11b^+^ macrophage and other non-T cells). **Figure 2E**: Total NK/NK-T cells versus 1D3-, FMC63-CD19CAR, and GFP transduced NK/NK-T cells (CD335^+^ NK / NKT cells). Flow gating strategy outlined in **Figure S4** a d data in **Table S3 and Table S4**.

To further assess *in vivo* transduced CAR-T cells from peripheral blood we used a panel of markers that included CD3, CD4, CD8, CD44, CD62L and CD127 [**Figure 4**]. A gating strategy for the use of these markers to define T cell immunophenotypes has been previously described [21,22], and an example of our implementation is illustrated in **Figure S5** and **Table S5**. At both timepoints tested (week 5 and week 8), we saw that CD8^+^ CAR-T cells were more prevalent than CD4^+^ CAR-T cells, and displayed mainly effector (CD44^high^, CD62L^low^ CD127^low^) or effector memory (CD44^high^, CD62L^low^ CD127^high^) phenotypes. Interestingly, these ID3 CAR-T cells proliferated sooner, and to a greater abundance than the FMC63 CAR-T cells, which did not become prominent until week 8. A small (<1%) but measurable population of GFP expressing CD4^+^ FOXP3^+^ regulator T cells was also observed at both timepoints. We conducted qPCR (Taqman) analysis to test for the presence of the CD19CAR transgene in additional tissues obtained 2 or 6 weeks after infusing mice with either of two different doses (2 x10^7^ or 2 x10^6^ IU) of the 1D3-CD19CAR-GFP virus. For the following experiments, the number of mice per treatment group was 7 (n=7). In **Figure 3**, error bars represent the standard deviation away from the calculated mean of each treatment group. From PBMC and spleen samples from treated mice we isolated DNA from three sorted cell populations: a T cell (CD3+) population, a B cell (CD20+) population, and an NK/NK-T and Macrophage (CD335+/CD11b+) population. We also isolated DNA from lung, ovary, bone marrow, and liver from these mice. At 6 weeks, mice treated with 2×10^6^ IU of virus showed a relative quantity of 132.7 +/-27.8 in PBMC-derived CD3^+^ T cells and 131.0 +/-35.4 in Spleen-derived CD3^+^ T cells, while mice treated with 2×10^7^ IU virus showed a relative quantity of 129.9 +/-34.8 in PBMC-derived CD3^+^ T cells and 147.1 +/-41.4 in Spleen isolated CD3^+^ T cells. Interestingly, at the earlier 2 week timepoint, qPCR analysis identified transduced T cells from PBMC and spleen in two of the six mice that were treated with the higher dose of 2×10^7^ IU of virus, suggesting that higher T cell transduction can promote the earlier appearance of a sizeable transduced T cell population, although only sporadically. None of the off target cell types or tissues were qPCR positive for the transgene at any of the doses or timepoints [**Figure 3**].

**Figure 3:**
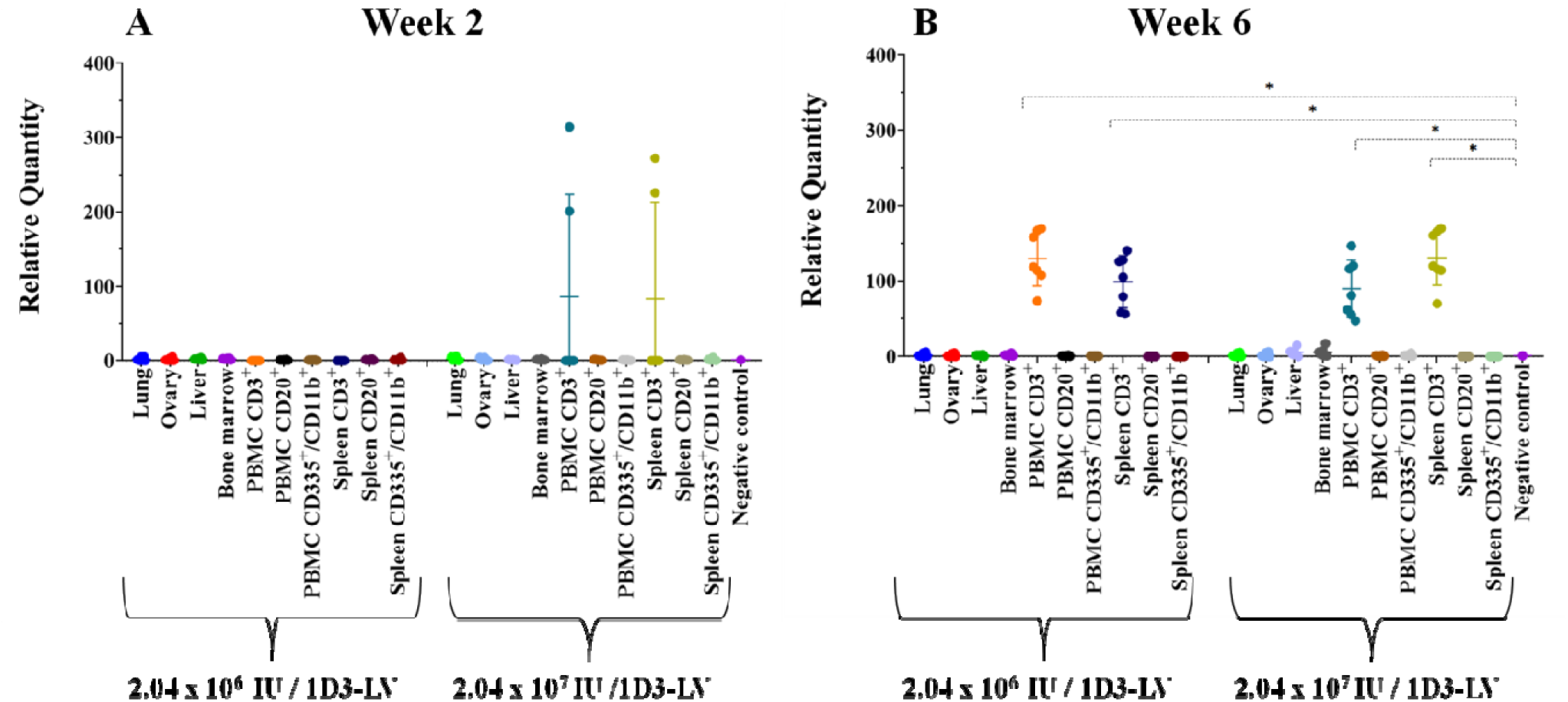
Quantitative PCR analysis of transgene positivity. DNA from CD3^+^, CD20^+^, and CD355^+^/11b^+^ cells, sorted from PBMC and spleen samples and from liver, lung, bone marrow, and the ovaries was obtained from female C57BL/6 mice. N equals 7 mice per group and the error bars represent the standard deviation from the mean value reported for each group. [**Figure 3A**] 2 weeks and [**Figure 3B**] 6 weeks after being treated with 2.04×10^6^ or 2.04×10^7^ IU of the 1D3-CD19CAR lentivirus, respectively. The 1D3-CD19CAR-GFP transgene were prominent only in the CD3^+^ T cell populations isolated from both PBMC and spleens 6 weeks posts 1D3-CD19CAR-LV treatment (**Figure 3B**. Orange 132.7 +/-27.77, dark blue 131.0 +/-35.42, cyan 129.9 +/-34.75, and mustard 147.1 +/-41.42). Relative quantity was calculated using the ΔΔCT calculation using the murine reference gene β-actin and unmodified C57BL/6 spleen cells as the reference control.We also see evidence of CD19CAR-T possession at week 2, in CD3^+^ T cells isolated from the spleen and PBMCs of two (2) mice which received 2.04 × 10^7^ IU of the 1D3-CD19CAR-LV (**Figure 3A**; cyan and mustard). **Figure 3A;** ANOVA ***p <0.001, **Figure 3B;** ANOVA***p <0.001. Using Dunnett’s post-hoc analysis, comparing PBMC and spleen T cells, 6 weeks after treatment with 2.04 × 10^6^ of the 1D3-CD19CAR-LV with the negative control, * p > 0.015.

**Figure 4:**
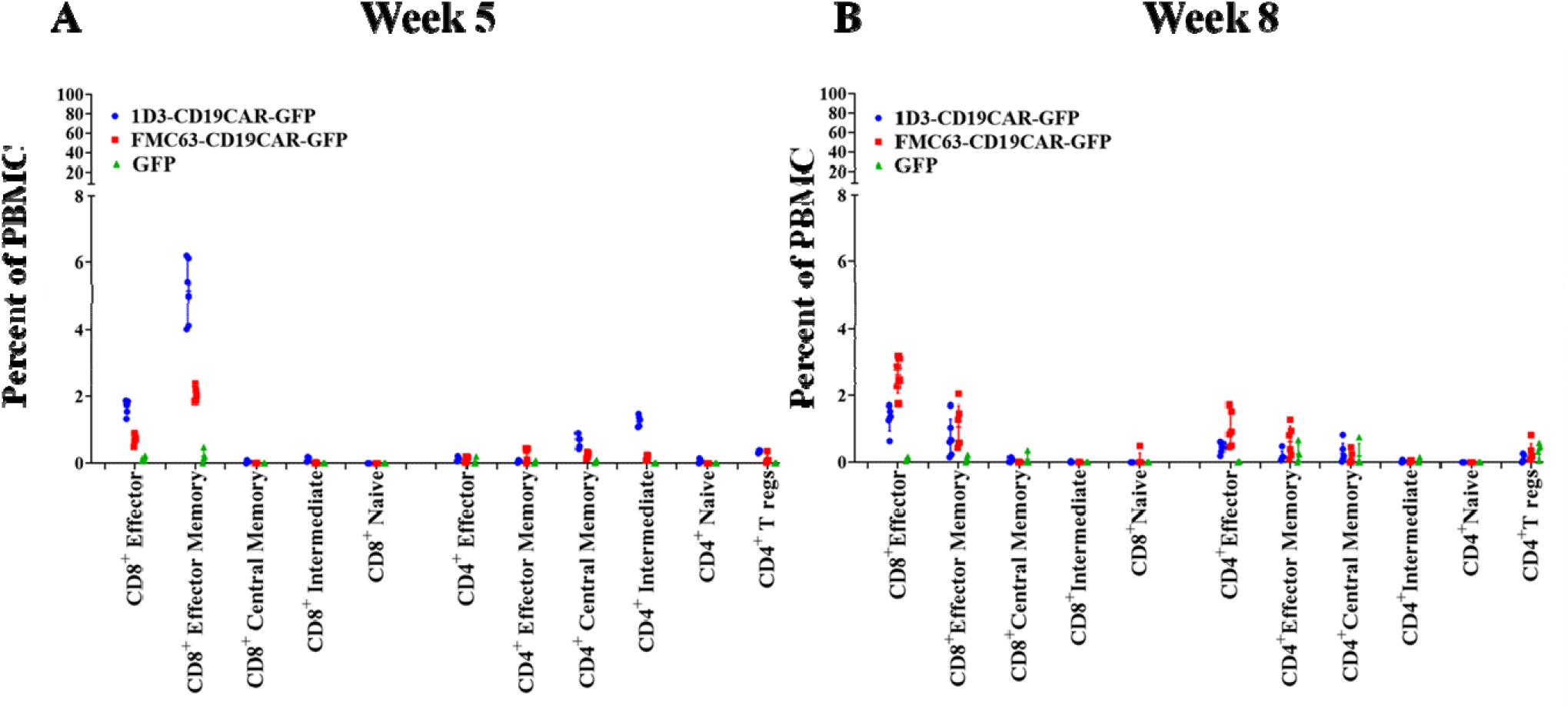
Immunophenotyping of 1D3-CD19CAR-GFP (blue, circle), FMC63-CD19 CAR-GFP (red, square) or control (GFP-only, green, triangle) transduced T cells in peripheral blood of treated mice. Values shown are the percentage of total CD3^+^, CD90.2^+^ T cells represented by each subtype (Effector = CD44^high^, CD62L^low^ CD127^low^ ; effector memory = CD44^high^, CD62L^low^ CD127^high^ ; central memory = CD44^high^, CD62L^high^ CD127^high^; intermediate phenotype = CD44^high^ CD62L^high^ CD127^low^; naive = CD44^low^, CD62L^high^ CD127^high^; regulatory = CD4^+^FOXP3^+^). N equals 8 mice per group and error bars represent the standard deviation from the mean value reported for each group. **Figure 4A**: At week 5 the 1D3-CD19CAR-GFP T cells were predominantly CD8^+^ effector memory T cells (5.1 +/-0.9% of total CD3^+^ T cells), effector T cells (1.7 +/-0.2% of total CD3^+^ T cells), and CD4^+^ intermediate T cells (1.3 +/-0.2% of total CD3^+^ T cells). At week 5 the majority of FMC63-CD19CAR-GFP T cells were CD8^+^ effector memory T cells (2.0 +/-0.2% of total CD3^+^ T cells) and effector T cells (0.7 +/-0.1% of total CD3^+^ T cells). **Figure 4B**: At week 8, the majority of 1D3-CD19CAR-GFP T cells were effector memory T cells (0.7 +/-0.6% of total CD3^+^ T cells) and CD8^+^ effector T cells (1.3+/-0.4% of total CD3^+^ T cells). The majority of FMC63-CD19CAR-GFP T cells at week 8 were CD8^+^ effector memory T cells (2.60 +/-0.538% of total CD3^+^ T cells) and effector T cells (1.1 +/-0.6% of total CD3^+^ T cells). GFP expressing CD4^+^ FOXP3^+^ regulator T cells were present at both time points but comprised of a small percentage (<1%) of total T cells. Flow gating strategy outlined in **Figure S5** and data in **Table S5**.

## Discussion

There have been no previously published reports of FMC63 crossreactivity, or that FMC63 is human specific. Zola et al.[19] initially characterized the FMC63 antibody as a mouse-human chimera with FMC63 mouse sequences for the VDJ (heavy) and VJ (light) chains with a human C region gene of IgG1, which was then tested for its ability to bind human CD19 compared to normal FMC63. Pietersz et al.[23] compared the mouse/human chimera FMC63 scFv for its ability to reduce human B cell tumours by 30% in scid/scid mice and found it similar to the mouse FMC63. Initial construction of the single chain Fv (scFv) of FMC63 [19] tested the scFv only against human cell lines. Construction of an anti-CD19 chimeric antigen receptor using this scFv [24]tested specifically against human PBMCs. Evolution of the anti-CD19 chimeric antigen receptor designs [25,26] tested against human CD19 tumours (NALM cell lines) but not against mouse cell lines. Evaluation of the chimeric antigen receptor-modified T cells [27]tested only against human cell lines and patient samples. None of these tested the effect of FMC63 against mouse CD19. The region of the FMC63 epitope (aa 155-172 [28]) has 75% amino acid identity and 83% amino acid similarity in human and mouse and structural prediction shows conserved conformation of this region (Figure S7).

Interestingly, murine T cells have previously been demonstrated to have lower transduction efficiencies, compared to human T cells and compared to other murine cell types [29–32], when transduced with HIV-1 based lentiviral vectors. The viral vector system we describe here avoids some of the known barriers of murine T cells. The entry block is avoided by the use of VSV-G pseudotyped virus. Reduced Tat activation is avoided entirely by our use of a Tat-independent transfer vector with a chimeric Rous Sarcoma virus LTR. Reduced LTR promoter activity is compensated by the use of an internal EF1α promoter driving the transgene. Some reduction in lentiviral integration may still be expected in murine T cells due to inefficient nuclear import of the pre-integration complex.[31] Empirically, however, we find that while our lentiviral vectors transduce murine T cells less efficiently than they transduce human T cells **[Figures S2]**, the level of transduction of murine T cells *in vitro* and *in vivo* is sufficient.

Quantitative PCR analysis at weeks 2 and 6 post initial injection, of T cells (CD3), B cells (CD20), NK/NK-T cells (CD335 and CD11b) as well as liver, lung, ovaries, and bone marrow tissue, showed that only the T cells had detectable CAR transgene. We did not look at the *in vivo* transduction of progenitor / stem cell types like CD34^+^ hematopoietic stem and progenitor cells or endothelial / vascular cell types. We expect that transduction events at our administered doses are below the limit of our detection, and that the transduction of the T cells are only detectable due to antigen driven CAR-T cell expansion. Indeed, only in the higher delivered viral dose, do we see any CAR transgene in the PBMC and spleen CD3+ T cells at the two week timepoint (**Figure 3**). Previous studies have reported transduction of hepatocytes by VSV-G pseudotyped lentiviral particles [33,34] when used in other settings. The substantially lower viral doses (2.04×10^7^ IU and 2.04×10^6^ IU per mouse) used in our study does not yield detectable levels of CAR transgene in other tissues. We would expect that B cells, NK/NK-T cells and macrophages, express low levels of LDL-R and thus the transduction of these cells by VSV-G lentivirus *in vivo* would be expected to be low [35] in accord with our lack of CAR transgene detection in these cells. However, given that other LDL-R bystander cell populations will act as viral sinks, and given that VSV-G lentiviruses are inactivated by human complement [36], optimal dosing regimens will need to be defined empirically.

Further implications to the transduction of bystander cells are unique to the setting of anti-CD19 CAR-T therapy. For example, expression of the CAR transgene in malignant B cells by inadvertent transduction of tumor blasts is potentially problematic, because the CAR can bind CD19 antigen on the same cell and block engagement by cytolytic CAR-T cells. This is a known but rare mechanism of relapse in conventional *ex vivo* protocols [37]. Another consideration is that while receptor transgene expression in CD8^+^ cytolytic T cells and CD4^+^ helper T cells is desirable,[38] expression in regulatory T cells may have a non-negligible inhibitory effect on CAR-T cell immunoreactivity over time. Utilizing modified promoters that drive CAR transgene expression in effector but not regulatory T cells is one strategy for overcoming this potential problem.

Insertional mutagenesis of integrating viral vectors led to highly regrettable negative outcomes in early gene therapy trials and severe setbacks to the gene therapy field. [39–44] All of these clinical trials involved *ex-vivo* gene transfer to hematopoietic stem cells using “first generation” gamma-retroviral vectors. Retrospective analysis has shown that the mechanism of leukemogenesis was oncogene overexpression driven by integration, in the vicinity of oncogenes, of viral vectors bearing strong promoters in their long terminal repeats. The advent of self-inactivating lentiviral vectors with transgene expression driven by mammalian (not viral) LTR promoters (as used in the present study) have mitigated oncogenic risk [45,46] and are now routinely used in different disease settings without incident, and no detrimental effects of insertional mutagenesis have been reported from any CAR-T clinical trial using retroviral vectors to date. Still, before clinical testing of *in vivo* CAR-T transduction could be considered, the risk of insertional mutagenesis must be fully mitigated.

The use of HIV-based lentivirus for gene transfer also raises concern about the theoretical risk of transmission of replication competent lentivirus (RCL). We use a modified 2^nd^ generation lentiviral vector system with the minimal viral components included in the 3^rd^ generation system split across three plasmids instead of four. These alterations abolish the potential for recombination events that can lead to formation of RCL in target cells. Testing of hundreds of cell therapy products generated with third generation vectors as well screening clinical trial participants after infusion of cell product failed to detect RCL, [47,48] demonstrating the safety of lentiviral gene therapy in this regard.

Clinical CAR-T protocols and other clinical IEC protocols typically include lymphodepleting chemotherapy with cyclophosphamide and/or fludarabine as pre-conditioning. While it would not be productive to perform complete lymphoablation prior to a viral infusion protocol (because endogenous T cells must be present for virus to transduce), mild preconditioning may be beneficial, if depletion leaves enough endogenous T cells in circulation to support viral transduction and conversion to CAR-Ts, while at the same time providing a niche for robust CAR-T expansion and engraftment [49]. This could be further tested, but may be unnecessary given our results show full ablation of endogenous B cells after lentiviral infusion **[Figure 2C]** without any pre-conditioning lymphodepletion.

In summary, anti-CD19 CAR-T therapy for B cell malignancies has shown clinical success, but a major limitation is the logistical complexity and high cost of manufacturing autologous cell products. This has led to increasing interest in industry and academia in universal (non-personalized) strategies. Here we report that intravenous infusion of replication-incompetent VSVG-pseudotyped lentiviral particles carrying an anti-CD19 CAR transgene with either an FMC63 or ID3 anti-CD19 binding domain can effectively transduce murine T cells, leading to CAR-T proliferation and subsequent CAR-T mediated B cell aplasia. Out study supports the notion that CAR-T cells expanded *in vivo* could be effective against B cell malignancies, which requires further study. This observation also supports this modality as a potential universal CAR-T intervention, that could have clinical relevance if further engineering is able to overcome the safety concerns and other limitations of systemic viral gene therapy.

## Materials and Methods

### Media

RPMI-1640 supplemented media (sRPMI-1640) consisted of 2 mM GlutaMAX (Invitrogen, 35050-061), 1 mM MEM non-essential amino acid (Invitrogen, 11140-050), 1 mM sodium pyruvate (Invitrogen, 11360-070), 10 mM HEPES (Invitrogen, 15630-808), 100 U/mL penicillin-streptomycin (Invitrogen, 15140-122), MycoZap prophylactic (Lonza, VZA-2031), 0.05mM 2-Beta Mercaptoethanol (Sigma, 60-42-2, 2-βME added to media for murine cell cultures only) and 10% heat-inactivated fetal bovine serum (Invitrogen, 12484-028). For expansion cultures IL-2 (StemCell, 78036.3 or Miltenyi Biotech, 130-097-745) was added to the sRPMI-1640 media. Cells were cultured in sRPMI-1640 with a final IL-2 concentration of 300IU/ml, unless otherwise stated. FACS media consisted of 1x D-PBS (Sigma, 59331C) with 2% HI-FCS. In house MACS separation buffer consisted of D-PBS with 2mM EDTA (ThermoFisher, 15575020) and 2% HI-FBS.

### Production of lentiviral gene transfer vectors

Our lentivector system is a modified 2^nd^ generation system that uses three separate plasmids to split the lentiviral genome as previously described.[50,51] The system is comprised of a packaging plasmid encoding gag, pol and rev, a VSV-G envelope expressing plasmid and a transfer plasmid encoding the transgene. Biosafety of the 2^nd^ generation design was improved by removal of tat from the packaging plasmid with the concurrent replacement of the transfer vector 5’ Long terminal repeat (LTR) with chimeric sequence for Tat-independent expression and a self-inactivating modified 3’ LTR.[52] These modifications serve to prevent the formation of replication competent virus. Synthetic transgene sequences encoding 1) GFP, 2) FMC63-CD19CAR-2A-GFP and 3) 1D3-CD19CAR-2A-GFP were manufactured (ATUM) containing the FMC63 (anti-human) or 1D3 (anti-mouse) anti-CD19 scFv with human co-stimulatory 4-1BB activation domains as a chimeric antigen receptor, subcloned into the transfer plasmid and verified by Sanger sequencing (Genewiz). Envelope (VSV-G) and packaging (GagPolRev) plasmids were also manufactured (ATUM). To manufacture virus particles, envelope, packaging, and transfer plasmids were co-transfected into the packaging cell line HEK-293T (clone 17; ATCC) using TransIT-LT1 (Mirus, MIR2305). Media containing the lentivirus was collected after transfection, filtered and then ultracentrifuged at 25,000 rpm or 106,750 xg, for 90 minutes at 4°C (Optima XE-90, Beckman-Coulter with SW 32 Ti swinging bucket rotor) to pellet the virus. Viral pellets were resuspended in 1x D-PBS and aliquoted for long term storage at -80C. Functional viral titers were established by infection of K562 cells (ATCC) and EL4 cells (ATCC) determined by transducing 5×10^4^ EL4 or K562 cells (ATCC) with 1, 2, 4, 8, 16, or 32 μL of concentrated viral supernatant. Performed in duplicate, cells were suspended in a final volume of 500μL of sRPMI-1640 media, in the 24-well format. Cells were then incubated for 72 hours, before being washed, centrifuged, and resuspended in FACS media. Acquisition was performed on the BD LSRFortessa cell analyser, and analysed using FlowJo and GraphPad Prism software version 8.0.0 for windows (GraphPad Software, California, USA) [**Figure S1]**.

### Sample acquisition and storage

Human PBMCs were isolated by Ficoll gradient purification as per manufactures protocol (Ficoll, GE-17-1440, GE Healthcare) from human peripheral blood leukapheresis pack (StemCell). 100-150μl of C57BL/6 mouse blood was collected in lithium heparin capillary tubes via tail vein blood sampling pre-treatment and then every week post-treatment for 8 weeks. The blood samples were diluted with 1 -1.5 ml of cold RPMI, which was overlayed with 4ml of Ficoll and C57BL/6 mouse PBMCs were isolated following the manufacturer’s protocol. C57BL/6 mouse PBMCs were isolated and analysed within 24 hours of isolation. PBMCs were resuspended in sRPMI-1640 at a concentration of 1×10^6^ cells per ml, placed into a T25 culture flask (156367, ThermoFisher), and rested overnight in the cell incubator at 37°C with 5% CO_2_ airflow before being used in any analysis. Human PBMCs and C57BL/6 mouse PBMCs not immediately used in any experiments were stored down by resuspending cells in freezing media (HI-FBS with 10% DMSO, Fisher Scientific, BP231-100) in aliquots of 5-10 x10^6^ cells per ml and frozen down to -80°C at a speed of 1°C/min prior to storage in liquid nitrogen.

The C57BL/6 mouse splenocytes used in the *in vitro* assays were acquired from naive or untreated C57BL/6 mice, aged 4-8 weeks old. All C57BL/6 mice were however processed the same way by first manually dissociating the spleens and filtering them through a 70 μm cell strainer (Corning, CLS431751) into a 50mL falcon tube, centrifuging 400xg for 10 min (Eppendorf Centrifuge 5810R, rotor A-4-62) at room temperature to pellet the cells. The splenocytes samples were resuspended in 5mL of ACK (Ammonium-Chloride-Potassium) Lysing Buffer (ThermoFisher), before incubating the cells on ice for no longer than 4 minutes. 10 mL of sRPMI-1640 media was added to the cells to neutralize the ACK buffer and wash the splenocytes. The splenocytes were then centrifuged at 400xg for 10 min (Eppendorf Centrifuge 5810R, rotor A-4-62) at room temperature, the supernatant was removed, and the splenocytes washed again before being resuspended in sRPMI-1640 media and cell numbers obtained using the Countess automated cell counter (ThermoFisher). Splenocytes were stored down by resuspending cells in freezing media (HI-FBS with 10% DMSO, Fisher Scientific, BP231-100) in aliquots of 5-10 x10^6^ cells per ml and frozen down to -80°C at a speed of 1°C/min prior to storage in liquid nitrogen.

The acquired bone marrow, ovaries, lung, and liver tissues were manually dissociated and filtered through a 70 μm cell strainer (Corning, CLS431751) into a 50mL falcon tube. Samples were then, washed with PBS and centrifuged at 400xg for 10 min (Eppendorf Centrifuge 5810R, rotor A-4-62) at room temperature to pellet the cells. The cells were counted and stored in RNA later (AM7021, ThermoFisher) following the manufactures protocols. DNA was extracted from these samples using the DNeasy Blood and Tissue kit (Qiagen, 69504) and following the manufacturer’s protocol.

### In vitro 1D3-and FMC63-CD19CAR-GFP modified T cell expansion and testing using C57BL/6 mouse splenocytes

C57BL/6 mouse splenocytes were removed from liquid nitrogen storage, thawed, and resuspended in sRPMI-1640 with 300 IU/ml of IL-2, and 30ng/ml of CD3 (clone 145-2C11, 100314, BioLegend) and 30ng/ml CD28 (clone 37.51, 102112, BioLegend) antibodies, at a concentration of 1×10^6^ cells/ml. The splenocytes were then treated with 1D3-, FMC63-CD19CAR-GFP, or GFP-only viral supernatant at a MOI of 5 (calculated based on titration data) and incubated for 72 hours. Following transduction, cells were washed by resuspending the cells in sRPMI-1640, centrifuging them at 400xg for 10 mins at room temperature (Eppendorf Centrifuge 5810R, rotor A-4-62), removing the supernatant, and repeating the process a further 2 times. The cells were resuspended in sRPMI-1640 media containing 300IU/ml of IL-2 at a concentration of 2.5×10^5^ cells per ml, with 1 ml being aliquoted per well, in the 6-well culture plate format. Irradiated autologous murine splenocytes (50Gy) were resuspended in sRPMI-1640 media containing 300IU/ml of IL-2 at a concentration of 2.5×10^7^/ml and 1 ml was added to the transduced splenocytes giving a final ratio of 1:100 transduced splenocytes to irradiated splenocytes. Every third day, 1ml of the culture media was removed and 1ml of fresh sRPMI-1640 media with 600IU/ml of IL-2 was added (giving a final concentration of 300IU/ml IL-2). On days 0, 14, 28, and 63, the cells were counted using the Countess automated cell counter (ThermoFisher) and an aliquot of 3×10^4^ cells were stained with fluorescent labeled antibodies and analyzed using flow cytometry [**Figure 1A** and **1B**].

In a 96-well plate format with a final volume of 200μl (sRPMI-1640), 1×10^5^ 1D3-, FMC63-CD19CAR, or GFP-transduced T cells from day 63 of the expansion were co-cultured for 5 hours with 1×10^5^ B cells isolated from another sample of C57BL/6 mouse splenocytes, using the Miltenyi MACS mouse CD19 B-cell isolation beads, following the manufacturer’s protocol and using a MACS separation buffer made in-house. At the end of the co-culture, the cells were stained with fluorescent labeled antibodies and analyzed using flow cytometry [**Figure 1C**]. Cells were washed with 500μl of D-PBS and centrifuged at 400xg for 10 minutes at room temperature (Eppendorf Centrifuge 5810R, rotor A-4-62). The supernatant was removed and the cell pellet resuspended in 50μl of D-PBS before being stained with CD20-PE (clone A1SB12, 12-0201-82, eBioscience), CD3-efluor 450 (clone 17A2, 48-0032-82, eBioscience), and the non-viable cell marker, Fixable Viability Stain 780 (FVS780) (565388, BD Biosciences). Cells and antibodies were incubated at 4ºC for 30 min, then washed by adding 2mL of FACS media to each tube and centrifuging for 10 minutes at 400xg at 4°C followed by removal of the supernatant before resuspension in 500μl of cold FACS media. Flow cytometry acquisition was performed using the BD LSRFortessa cell analyser with BD FACSDiva software. The analysis was performed using FlowJo software, version 10 for windows and GraphPad Prism software version 8.0.0 for windows (GraphPad Software, California, USA) [**Figure S3** and **Table S1**].

### In vitro 1D3-and FMC63-CD19CAR-GFP modified mouse T cells co-culture with A20 cells

C57BL/6 mouse splenocytes were transduced with 1D3-, FMC63-CD19CAR-GFP, or GFP-only viral supernatant, sorted, and expanded as previously mentioned. The murine, CD19 expressing, A20 cell line (ATCC) was used as the target cells in a co-culture with the sorted and expanded 1D3-, FMC63-CD19CAR-GFP, or GFP-only expressing murine T cells. In order to verify the ability for both the 1D3- and FMC63-CD19CAR to target the mouse CD19 antigen, the A20 cells were incubated with 10, 1, 0.1, 0.001, 0 μg/ml of either the 1D3-CD19 blocking antibody (152402, BioLegend) or the FMC63-CD19 blocking antibody (NBP2-52716, Novus Biologicals). In the 96-well plate format with a final volume of 200μl (sRPMI-1640) the A20 cells were then co-cultured the 1D3-, FMC63-CD19CAR-GFP, or GFP-only expressing murine T cells at an effector to target of 4:1(2×10^5^:5×10^4^), for 4 hours. Cells were then washed and stained with CD20-PE, CD3-efluor 450, and the non-viable cell marker, Fixable Viability Stain 780 (FVS780) as previously described [**Figure S3** and **Table S2**]. Flow cytometry acquisition was performed using the BD LSRFortessa cell analyser with BD FACSDiva software. The analysis was performed using FlowJo software, version 10 for windows and GraphPad Prism software version 8.0.0 for windows (GraphPad Software, California, USA) [**Figure 1D**].

### In vivo testing

All animal experiments were assessed and approved by the University of British Columbia’s Animal Care Committee under ethics certificate #A17-0107. For the *in vivo* testing, C57BL/6 mice, aged 4 to 8 weeks old, were obtained from the BC Cancer Research Centre’s Animal Resource Centre. The experimental mice received one tail vein infusion on day 1 which involved the intravenous injection of 200μl of D-PBS containing 3.6-4×10^6^ total infectious units of either 1D3-, FMC63-CD19CAR-GFP, or GFP-only. For subsequent dose response *in vivo* experiments, mice received an intravenous injection of 200μl of D-PBS containing either 2.04×10^6^ or 2.04×10^7^ total infectious units of 1D3-CD19CAR-GFP. Mice were monitored closely for the first 72 hours, and then once a day until the experimental endpoint. The mice were also weighed periodically and blood collected in lithium heparin capillary tubes via tail vein sampling as previously described. At the experimental endpoint, the mice were humanly euthanized and spleen and blood tissue collected and processed as previously described. The cells were stained with fluorescent labeled antibodies and analyzed using flow cytometry [**Figure 2** and **Figure 4**]. DNA from the cells of the various tissues was collected and the samples analyzed for CD19CAR-GFP possession using qPCR [**Figure 3**].

For flow cytometry acquisition and analysis, 4×10^5^ cells were stained with CD3-efluor 450 (clone 17A2, 48-0032-82, eBioscience), CD90.2-AF700 (clone 30-H12, 105319, BioLegend), CD20-PE (clone A1SB12, 12-0201-82, eBioscience), CD45R/B220-APC-CY7 (clone RA3-6B2, 103223, BioLegend), CD335-APC (clone 29A1.4, 137608, BioLegend), and CD11B-BV605 (clone M1/79, 101237, BioLegend) conjugated flow antibodies. Cells and antibodies were incubated at 4ºC for 30 min, then washed by adding 2mL of FACS media to each tube and centrifuging for 10 minutes at 400xg at 4°C, discarding the supernatant before resuspension in 500μl of cold FACS media. Acquisition was performed on the BD LSRFortessa cell analyser, and analysed using FlowJo and GraphPad Prism software version 8.0.0 for windows (GraphPad Software, California, USA) [**Figure S4, Table S3** and **Table S4**].

In the immunophenotyping analysis [**Figure 4**], up to 5×10^4^ cells were stained with CD3-efluor 450 (clone 17A2, 48-0032-82, eBioscience), CD4-PE (clone GK1.5, 100408, BioLegend), CD8-BUV395 (clone QA17A07, 155006, BioLegend), CD44-alexa fluor 700 (clone IM7, 103026, BioLegend), CD62L-PerCP-CY5.5 (clone MEL-14, 104432, BioLegend), CD127-APC-CY7 (clone A7R34, 135040, BioLegend), CD25-PE-CF594 (clone PC61, 562694, BD Biosciences), and FOXP3-APC (clone FJK-16s, 563786, BD Biosciences) and incubated at 4ºC for 30 min, as previously mentioned. The cells were then fixed, by incubating the cells in up to 200μL of fixation buffer (420801, BioLegend) at 4ºC for 30 min. The cells were than washed by adding 2mL of FACS media and centrifuging for 10 minutes at 400xg at 4°C, and resuspended in 1x Intracellular Staining Perm Wash Buffer (10x stock solution, diluted to 1x with ddH_2_O, 420801, BioLegend) at 4ºC for 30 min, before being stained with FOXP3-APC (at 4ºC for 30 min). Again cells were washed by adding 2mL of FACS media and centrifuging for 10 minutes at 400xg at 4°C, discarding the supernatant before resuspension in 500μl of cold FACS media. Acquisition was performed on the BD LSRFortessa cell analyser, and analysed using FlowJo and GraphPad Prism software version 8.0.0 for windows (GraphPad Software, California, USA) [**Figure S5 and Table S5**]. For all flow analyses, we used the CD20-PE antibody as a B cell marker in place of a CD19 antibody [**Figure 1** and **Figure 2**] to avoid any possible interference between anti-CD19 antibody and anti-CD19 CARs in these mixed cell populations.

### Quantitative PCR of DNA isolated from mouse tissues

Spleens and PBMC from 2 and 6 weeks post 1D3-CD19CAR-LV treatment were sorted into CD3, CD20, and CD335/CD11b expressing cell types using BD FACSAria and BD FACSDiva software [**Figure S4**]. DNA was then extracted from the spleen and PBMC sorted cell subtypes, along with the DNA from liver, ovaries, lung, and bone marrow tissues (stored in RNA later) was collected using the DNeasy Blood and Tissue kit (Qiagen, 69504) following the manufacturer’s protocol. The DNA concentrations were read using NanoDrop 8000 Spectrophotometer (ND-8000-GL, ThermoFisher) by following the manufacturer’s protocol. The DNA concentration was normalized to a concentration of 19-21ng/μl. To detect the presence of the CD19CAR gene, using a Taqman primer probe set which amplifies DNA from the co-stimulatory and activation domains of the 1D3-CD19CAR-T transgene. For every 2μl of template DNA, 0.5μl of the CD19CAR [44] and murine-β actin Taqman assay sets (4352341, ThermoFisher), 5μl of Taqman Fast Advanced 364 Master Mix (4444557, Invitrogen), brought to a final volume of 10μl, with ultrapure water (10977023, Invitrogen). Thermocycling was done using the QuantStudio 6 Flex System, in the 384-Well format, using the comparative Ct (ΔΔCt) set up. Thermocycling conditions: one cycle at 50°C for 2 min; 1 cycle at 95°C for 2 min, 368 then 95°C for 2 sec and 60°C for 20 sec for 40 cycles, capturing data at the end of every cycle. For the analysis, the mouse β-Actin gene was used as the endogenous control and an *in vitro* transduced spleen sample, positive for 1D3-CD19CAR-GFP possession, was used as a positive control. Spleen cells from an untreated mouse was used as the reference control for the ΔΔCT calculation required to obtain a relative quantity.

Thermocycling was done using the QuantStudio 6 Flex System, in the 384-Well format, using the comparative Ct (ΔΔCt) set up. Thermocycling conditions: one cycle at 50°C for 2 min; 1 cycle at 95°C for 2 min, then 95°C for 3 sec and 60°C for 30 sec for 40 cycles, capturing data at the end of every cycle. For the analysis, the mouse β-Actin gene was used as the endogenous control and the CD3^+^ PBMC and Spleen from PBS treated mouse from week 1 and week 8 were used as the reference control, as these samples showed the lowest relative quantity, as they were treated with PBS.

## Author Contributions and Notes

R.A.H, C.M.R and E.Y. designed research, C.M.R, E.Y, L.D., and D.J.W. performed research, C.M.R and S.D.B. analyzed data; all authors prepared and edited the manuscript.

The authors declare no conflict of interest.

## Acknowledgments

This research was supported by the British Columbia Cancer Foundation, the Leon Judah Blackmore Foundation and the BioCanRx network.

## Supplementary Materials

**Figure S1:**
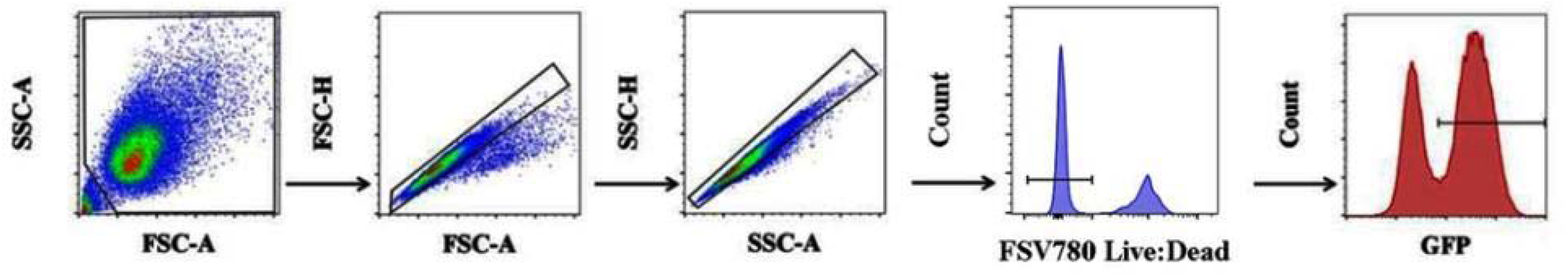
Viral supernatant titration flow gating protocol. After debris was eliminated by FSC-A vs. SSC-A gating, cell doublets and clumps were then eliminated by FSC-H vs. FSC-A gating followed by SSC-A vs. SSC-H gating. The live cells were determined by gating on the FVS780 negative cells and then the GFP positive cells were determined to establish the percentage of GFP expressing cells in order to measure transduction efficiency. The flow collected data was analysed using Flow Jo v10, for windows and GraphPad Prism software version 8.0.0 for windows (GraphPad Software, California, USA).

**Figure S2:**
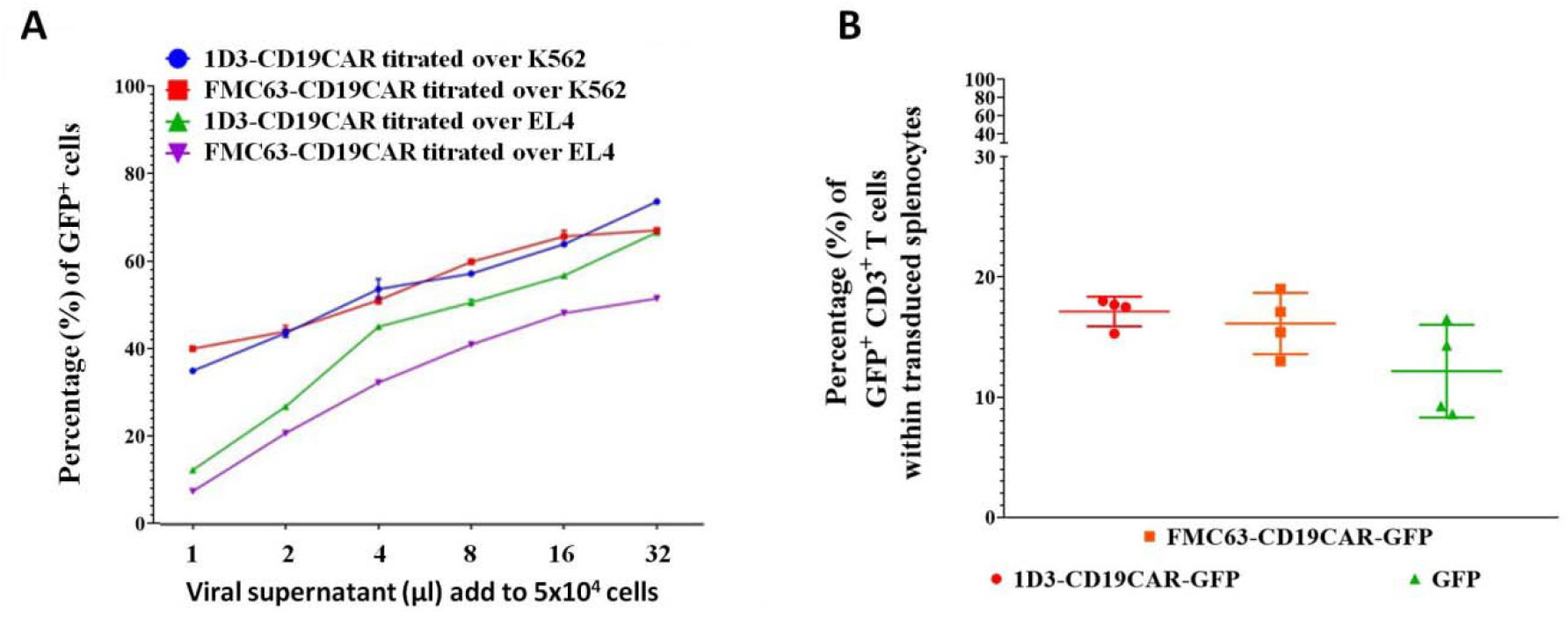
1D3-CD19CAR-GFP, FMC63-CD19CAR-GFP, and GFP lentivirus transduction efficiency. Figure S2A. Titration of 1D3-CD19CAR-GFP and FMC63-CD19CAR-GFP viral supernatant titration over EL4 (green and purple, respectively) and K562 (red and blue, respectively) cell lines. While the transduction efficiency appears significantly better in the human K562s (p < 0.001 ANOVA), the transduction of a murine cell line, EL4, indicates this lentivirus is able to transduce murine cells, albeit at a lower frequency. **Figure S2B**: 2.5×10^5^ splenocytes were transduced with 1D3-CD19CAR-GFP, FMC63-CD19CAR-GFP, and GFP lentivirus at an MOI of 5 (calculated based on Figure S2A). The percentage (%) of GFP^+^, CD3+ T cells within the whole splenocyte sample was determined by assessing 3.0×10^4^ cells via flow cytometry 72 hours after transduction with virus. 1D3-CD19CAR-GFP had a transduction efficiency of 17.1% (+/-1.23%), FMC63-CD19CAR-GFP 16.3% (+/-2.6%), and GFP 12.2% (+/-3.85%).

**Figure S3:**
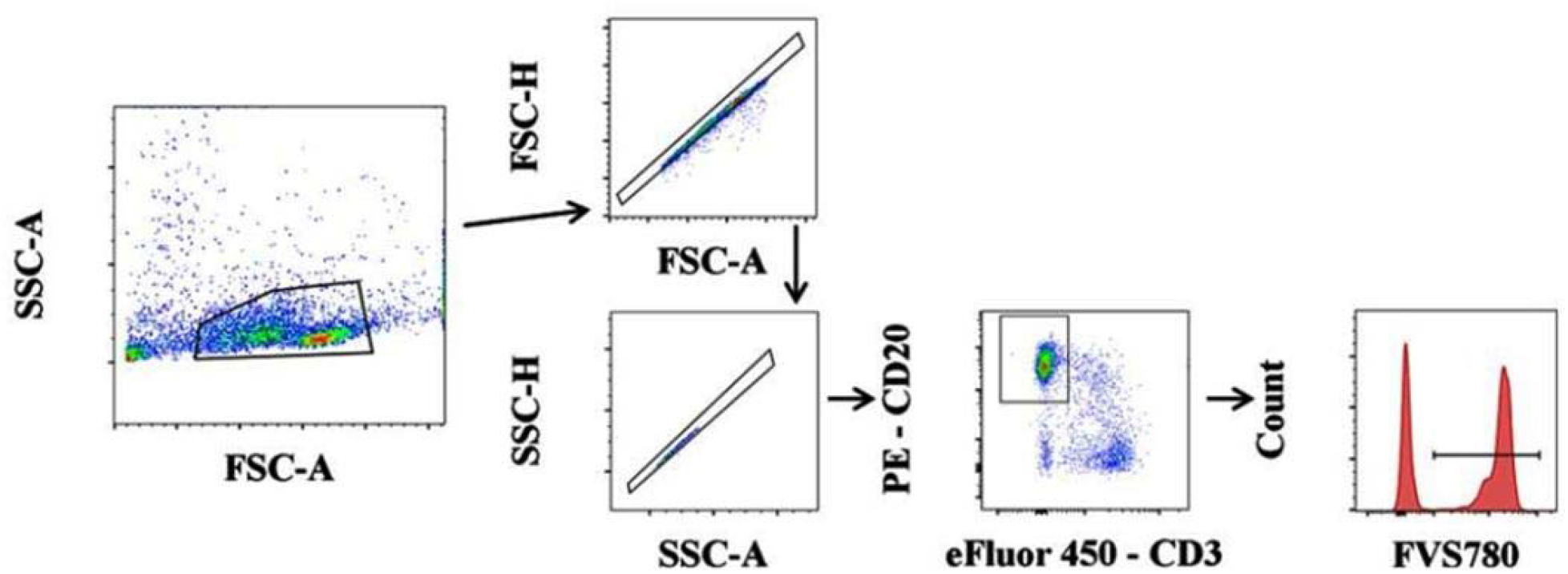
*In vitro* 1D3-or FMC63-CD19CAR-GFP T cell cytolytic activity using C57BL/6 splenocytes and A20 cells blocked with serially diluted concentrations of CD19 blocking antibodies. Debris, cell doublets, and clumps were eliminated by FCS-A vs SSC-A gating followed by FSC-A vs. FSC-H and SSC-A vs. SSC-H gating. T and B cells were then identified by CD3 vs. CD20 gating. CD20 positive cells were selected for and gated on FVS-780 to obtain the percentage of live vs. dead target cells. The flow collected data was analysed using Flow Jo v10, for windows and GraphPad Prism software version 8.0.0 for windows (GraphPad Software, California, USA) [**Figure 1**]

**Table S1:** Cell numbers for the *In vitro* 1D3- or FMC63-CD19CAR-GFP T cell cytolytic activity using C57BL/6 splenocytes data showed in **Figure 1C** and **Figure S3**

|  | **1D3-CD19CAR-GFP** |  | **FMC63-CD19CAR-GFP** |  |
| --- | --- | --- | --- | --- |
|  | <u>Mean cell count</u> | <u>SD</u> | <u>Mean cell count</u> | <u>SD</u> |
| CD19CAR-GFP | 12387 | 845.6 | 9263 | 540.6 |
| GFP | 167 | 120.1 | 400 | 658.6 |

**Table S2:** Cell numbers for the *In vitro* 1D3- or FMC63-CD19CAR-GFP T cell cytolytic activity using A20 cells blocked with serially diluted concentrations of CD19 blocking antibodies data showed in **Figure 1D** and **Figure S3**.

| **Blocking Ab (µg/ml)** | **1D3-CD19CAR-GFP** |  | **FMC63-CD19CAR-GFP** |  | **GFP** |  |
| --- | --- | --- | --- | --- | --- | --- |
|  | <u>Mean cell count</u> | <u>SD</u> | <u>Mean cell count</u> | <u>SD</u> | <u>Mean cell count</u> | <u>SD</u> |
| 0.00 | 19400 | 353.6 | 20525 | 459.6 | 9100 | 353.6 |
| 0.01 | 17000 | 636.4 | 17425 | 247.5 | 9025 | 106.1 |
| 0.10 | 14050 | 212.1 | 14100 | 495.0 | 9100 | 353.6 |
| 1.00 | 11400 | 282.8 | 10950 | 565.7 | 9025 | 106.1 |
| 10.0 | 9850 | 141.4 | 10400 | 282.8 | 9100 | 353.64 |

**Figure S4:**
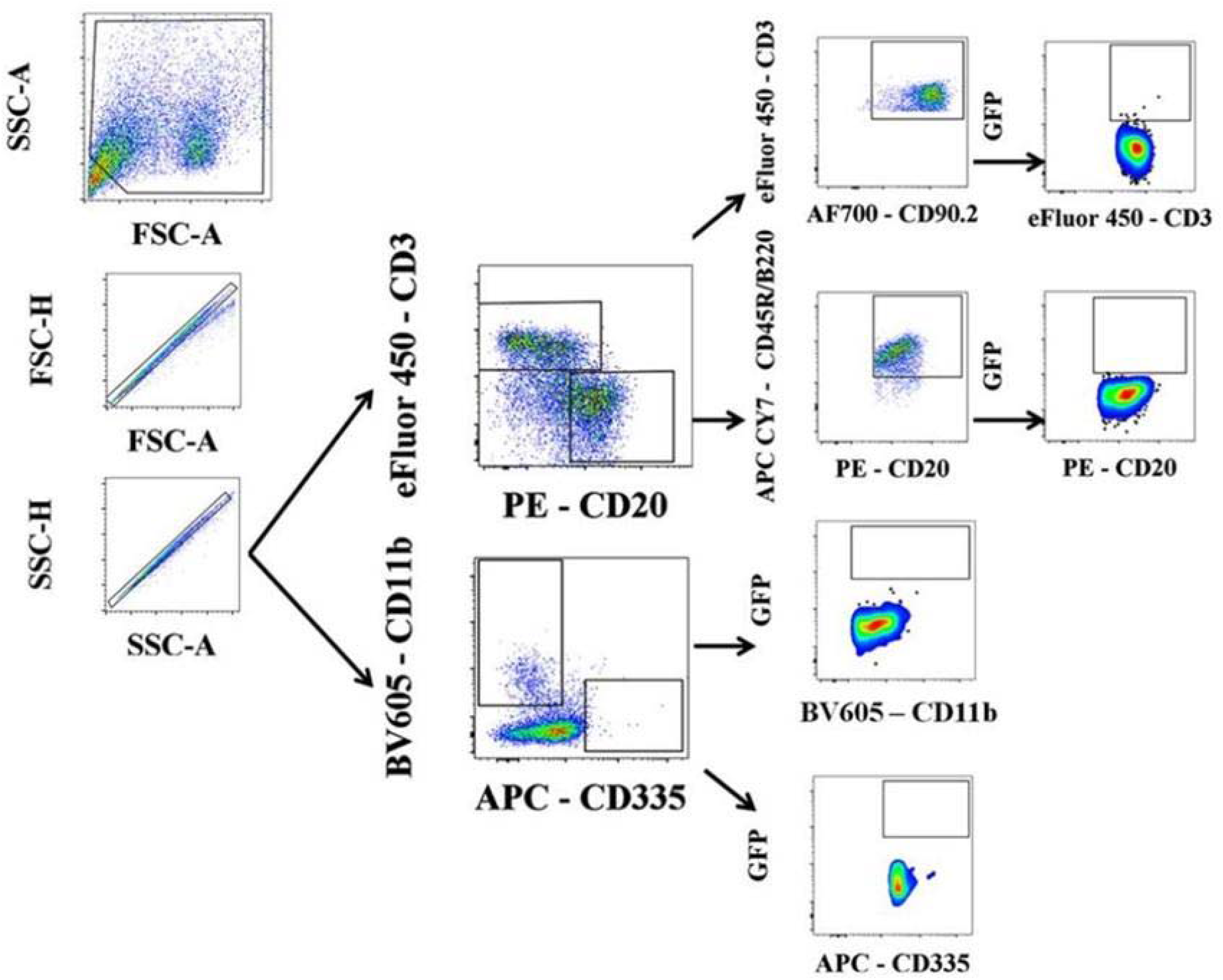
Flow gating protocol for the analysis of mouse blood pre- and post-treatment with lentiviruses. Debris was eliminated by FSC-A vs. SSC-A gating, cell doublets and clumps were then eliminated by FSC-H vs. FSC-A gating followed by SSC-A vs. SSC-H gating. T and B cells were then identified based on expression of CD3 or CD20 and NK cells and monocytes based on the expression of CD335 or CD11b, respectively. The CD3 positive cells were then gated for expression of CD90.2, and these CD3 and CD90.2 positive cells were gated for GFP expression (the CD19CAR reference gene marker). CD20 positive cells were gated for CD45R/B220 expression, and the CD20 and CD45R/B220 positive cells were then gated for GFP expression. NK cells (CD335) or monocytes (CD11b) were individually gated for GFP expression. The flow collected data was analysed using Flow Jo v10, for windows and GraphPad Prism software version 8.0.0 for windows (GraphPad Software, California, USA) [**Figure 2**]

**Table S3:** Cell numbers for B and T cells from flow cytometry data collected in the *in vivo* experiments, shown in **Figure 2** and **Figure 3** and flow gating protocol shown in **Figure S5**

|  | Week | GFP+ B cells |  | B cells |  | GFP+ T cells |  | T cells |  |
| --- | --- | --- | --- | --- | --- | --- | --- | --- | --- |
|  |  | Mean cell count | SD | Mean cell count | SD | Mean cell count | SD | Mean cell count | SD |
| **1D3-CD19CAR-GFP** | 0 | 4 | 6.13 | 15943 | 101.8 | 6 | 5.8 | 19214 | 140.6 |
|  | 1 | 1 | 3.6 | 20479 | 1220.3 | 84 | 42.48 | 18043 | 1511.7 |
|  | 2 | 2 | 5.1 | 19128 | 1382.0 | 53 | 17.68 | 19943 | 1567.8 |
|  | 3 | 23 | 39.5 | 14807 | 474.7 | 2421 | 255.6 | 14807 | 474.7 |
|  | 4 | 0 | 0 | 12043 | 4772.6 | 2973 | 711.8 | 17100 | 2409.7 |
|  | 5 | 0 | 0 | 144 | 32.3 | 6707 | 290.7 | 15764 | 536.0 |
|  | 6 | 0 | 0 | 20 | 13.3 | 2166 | 124.3 | 21329 | 722.2 |
|  | 7 | 0 | 0 | 294 | 77.2 | 3944 | 1446.1 | 20014 | 3161.9 |
|  | 8 | 0 | 0 | 275 | 96.9 | 3346 | 2163.8 | 17329 | 3298.9 |
| **FMC63-CD19CAR-GFP** | 0 | 0 | 0 | 14883 | 169.3 | 17 | 14.1 | 19425 | 181.0 |
|  | 1 | 12 | 18.83 | 20767 | 1423.6 | 67 | 19.7 | 18183 | 1702.3 |
|  | 2 | 3 | 7.3 | 19933 | 2273.0 | 75 | 18.4 | 18317 | 2166.7 |
|  | 3 | 10 | 24.5 | 22425 | 1221.0 | 775 | 84.7 | 16575 | 504.7 |
|  | 4 | 0 | 0 | 9833 | 1792.3 | 3061 | 655.6 | 14433 | 2556.3 |
|  | 5 | 0 | 0 | 281 | 281.1 | 3890 | 378.9 | 14925 | 503.7 |
|  | 6 | 0 | 0 | 46 | 20.4 | 2099 | 158.9 | 16133 | 503.7 |
|  | 7 | 0 | 0 | 429 | 56.3 | 4239 | 1345.0 | 20925 | 3533.8 |
|  | 8 | 0 | 0 | 255 | 49.2 | 1880 | 575.0 | 19700 | 1882.0 |
| **GFP** | 0 | 2 | 5.0 | 15343 | 395.2 | 14 | 22.3 | 19357 | 185.8 |
|  | 1 | 3 | 5.5 | 19157 | 1745.1 | 66 | 36.8 | 19671 | 2002.7 |
|  | 2 | 11 | 12.4 | 19550 | 1869.5 | 59 | 22.8 | 19150 | 2076.7 |
|  | 3 | 0 | 0 | 22279 | 1749.0 | 29 | 18.1 | 17179 | 1689.6 |
|  | 4 | 0 | 0 | 22679 | 2342.3 | 15 | 18.8 | 17614 | 1581.1 |
|  | 5 | 0 | 0 | 21571 | 2655.3 | 35 | 35.8 | 17850 | 798.4 |
|  | 6 | 0 | 0 | 22928 | 2501.0 | 116 | 42.4 | 20364 | 2216.0 |
|  | 7 | 15 | 22.8 | 17776 | 3410.6 | 33 | 11.7 | 21636 | 34 |
|  | 8 | 13 | 5.8 | 15157 | 3602.4 | 129 | 66.1 | 16471 | 3576.3 |

**Table S4:** Cell numbers for CD335^+^ and CD11b^+^ cells from flow cytometry data collected in the *in vivo* experiments, shown in **Figure 2** and **Figure 3** and flow gating protocol shown in **Figure S5**

|  | Week | GFP+ CD355 cells |  | CD355 cells |  | GFP+ CD11b cells |  | CD11b cells |  |
| --- | --- | --- | --- | --- | --- | --- | --- | --- | --- |
|  |  | Mean cell count | SD | Mean cell count | SD | Mean cell count | SD | Mean cell count | SD |
| **1D3-CD19CAR-GFP** | 0 | 0 | 0 | 1172 | 70.4 | 0 | 0 | 4392 | 62.4 |
|  | 1 | 0 | 0 | 1661 | 102.7 | 0 | 0 | 4482 | 369.6 |
|  | 2 | 0 | 0 | 1186 | 250.5 | 148 | 99.5 | 2402 | 216.2 |
|  | 3 | 0 | 0 | 614 | 275.9 | 0 | 0 | 2487 | 537.2 |
|  | 4 | 14 | 22.5 | 1104 | 407.8 | 14 | 22.5 | 2550 | 247.2 |
|  | 5 | 0 | 0 | 1228 | 418.5 | 0 | 0 | 2441 | 261.2 |
|  | 6 | 369 | 236.6 | 1076 | 50.2 | 44 | 117.2 | 556 | 24.7 |
|  | 7 | 91 | 241.9 | 797 | 194.8 | 213 | 134.0 | 3769 | 1415.9 |
|  | 8 | 140 | 370.4 | 659 | 211.9 | 496 | 343.6 | 1098 | 747.0 |
| **FMC63-CD19CAR-GFP** | 0 | 0 | 0 | 1118 | 37.0 | 0 | 0 | 4696 | 64.1 |
|  | 1 | 0 | 0 | 1638 | 73.9 | 0 | 0 | 4075 | 375.5 |
|  | 2 | 0 | 0 | 1324 | 385.6 | 33 | 53.3 | 2704 | 226.0 |
|  | 3 | 0 | 0 | 1038 | 402.3 | 0 | 0 | 1303 | 351.6 |
|  | 4 | 33 | 8.3 | 917 | 272.4 | 35 | 8.3 | 1772 | 110.4 |
|  | 5 | 0 | 0 | 906 | 244.0 | 0 | 0 | 1887 | 139.5 |
|  | 6 | 168 | 119.9 | 1371 | 80.8 | 0 | 0 | 699 | 34.6 |
|  | 7 | 0 | 0 | 707 | 104.1 | 155 | 136.0 | 2597 | 1541.6 |
|  | 8 | 34 | 83.7 | 612 | 80.4 | 366 | 346.4 | 479 | 46.4 |
| **GFP** | 0 | 0 | 0 | 1163 | 63.4 | 0 | 0 | 4639 | 105.9 |
|  | 1 | 0 | 0 | 1694 | 69.7 | 0 | 0 | 4093 | 273.0 |
|  | 2 | 0 | 0 | 1318 | 279.1 | 27 | 45.3 | 2512 | 328.8 |
|  | 3 | 0 | 0 | 1246 | 413.2 | 26 | 32.7 | 2298 | 710.5 |
|  | 4 | 0 | 0 | 1662 | 882.4 | 0 | 0 | 2114 | 623.5 |
|  | 5 | 0 | 0 | 1737 | 904.0 | 0 | 0 | 2234 | 675.3 |
|  | 6 | 0 | 0 | 1663 | 218.2 | 0 | 0 | 477 | 136.9 |
|  | 7 | 0 | 0 | 558 | 232.3 | 0 | 0 | 1969 | 1886.9 |
|  | 8 | 0 | 0 | 895 | 207.6 | 0 | 0 | 960 | 270.3 |

**Figure S5:**
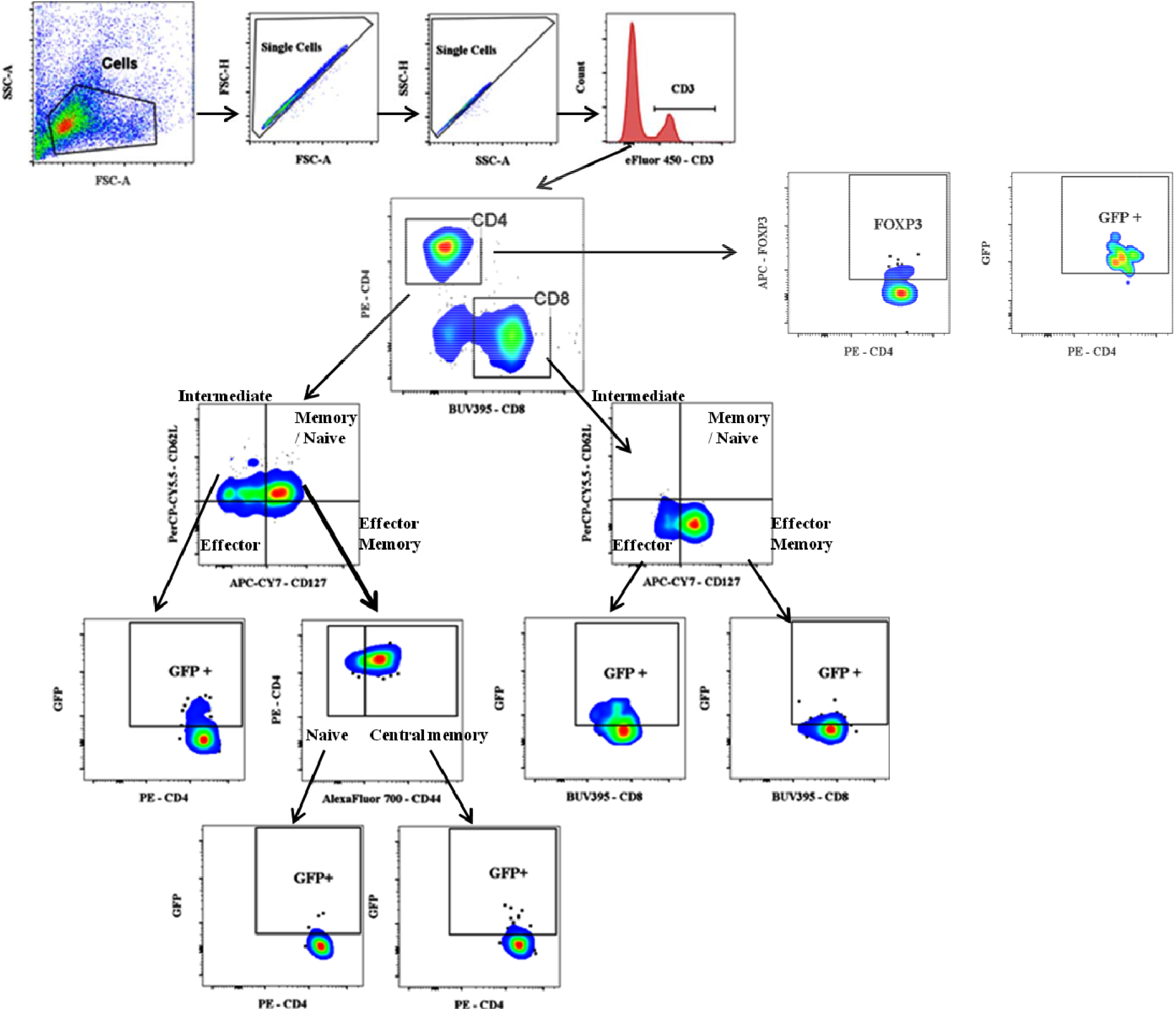
Flow gating protocol for the immunophenotyping analysis of murine T cells 5 and 8 weeks after lentiviral treatment. All cells were gated for in the FSC-A vs. SSC-A plot, cell doublets and clumps were then eliminated by FSC-H vs. FSC-A gating followed by SSC-A vs. SSC-H gating. T cells were then identified based on expression of CD3. The CD3 positive cells were then gated for expression of CD4 and CD8, and these CD3^+^ and CD8^+^ or CD4^+^positive cells were gated for CD127 and CD62L expression, allowing us to differentiate Central memory and naive cells, effector, and effector memory T cells. The Central Memory and Naive cells were then identified by their CD44 expression profile. Each of these subtypes were then looked at for GFP expression (the CD19CAR reference gene marker) indicated by the example plots above showing the positive expression of GFP. For regulatory T cells we plotted the CD4^+^ against the FOXP3, and then looked the number which expressed GFP. The flow collected data was analysed using Flow Jo v10, for windows and GraphPad Prism software version 8.0.0 for windows (GraphPad Software, California, USA) [**Figure 4**].

**Table S5:** Cell numbers for Immunophenotyping of 1D3-CD19CAR-GFP, FMC63-CD19 CAR-GFP, and GFP-only transduced T cells in peripheral blood of treated mice at week 5 and week 8 post treatment shown in **Figure 4** and flow gating protocol shown in **Figure S5**

|  |  | 1D3-CD19CAR-GFP |  | FMC63-CD19CAR-GFP |  | GFP |  |
| --- | --- | --- | --- | --- | --- | --- | --- |
|  |  | Mean cell count | SD | Mean cell count | SD | Mean cell count | SD |
| Week 5 | CD8<sup>+</sup>Effector | 1701 | 203.1 | 733 | 138.9 | 139 | 75.4 |
|  | CD8<sup>+</sup>Effector Memory | 5135 | 938.0 | 2029 | 205.0 | 215 | 203.4 |
|  | CD8<sup>+</sup>Central Memory | 42 | 34.6 | 8 | 8.5 | 2 | 4.25 |
|  | CD8<sup>+</sup>Intermediate | 109 | 58.9 | 7 | 13.6 | 0 | 0.0 |
|  | CD8<sup>+</sup>Naive | 0 | 0.0 | 0 | 0.0 | 0 | 0.0 |
|  | CD4<sup>+</sup>Effector | 121 | 61.5 | 131 | 77.3 | 53 | 105.0 |
|  | CD4<sup>+</sup>Effector Memory | 62 | 26.83 | 311 | 196.6 | 29 | 38.4 |
|  | CD4<sup>+</sup>Central Memory | 682 | 172.93 | 236 | 83.1 | 45 | 41.6 |
|  | CD4<sup>+</sup>Intermediate | 1262 | 164.3 | 154 | 76.5 | 3 | 6.4 |
|  | CD4<sup>+</sup>Naive | 55 | 57.2 | 0 | 0 | 0 | 0 |
|  | CD4<sup>+</sup>T reg | 355 | 32.7 | 118 | 128.3 | 3 | 6.4 |
| Week 8 | CD8<sup>+</sup>Effector | 1320 | 363.2 | 2604 | 538.3 | 110 | 52.5 |
|  | CD8<sup>+</sup>Effector Memory | 742 | 573.2 | 1075 | 637.5 | 98 | 99.0 |
|  | CD8<sup>+</sup>Central Memory | 71 | 52.2 | 3 | 6.8 | 91 | 182.5 |
|  | CD8<sup>+</sup>Intermediate | 10 | 16.3 | 4 | 4.8 | 4 | 7.25 |
|  | CD8<sup>+</sup>Naive | 2 | 2.5 | 83 | 204.1 | 0 | 0 |
|  | CD4<sup>+</sup>Effector | 460 | 169.3 | 1003 | 523.6 | 16 | 11.7 |
|  | CD4<sup>+</sup>Effector Memory | 186 | 155.4 | 634 | 465.3 | 252 | 304.0 |
|  | CD4<sup>+</sup>Central Memory | 270 | 309.2 | 127 | 190.0 | 191 | 382.5 |
|  | CD4<sup>+</sup>Intermediate | 25 | 31.0 | 38 | 25.1 | 39 | 77.5 |
|  | CD4<sup>+</sup>Naive | 1 | 2.2 | 0 | 0.0 | 0 | 0.0 |
|  | CD4<sup>+</sup>T reg | 102 | 114.8 | 289 | 283.3 | 362 | 234.8 |

**Figure S6:**
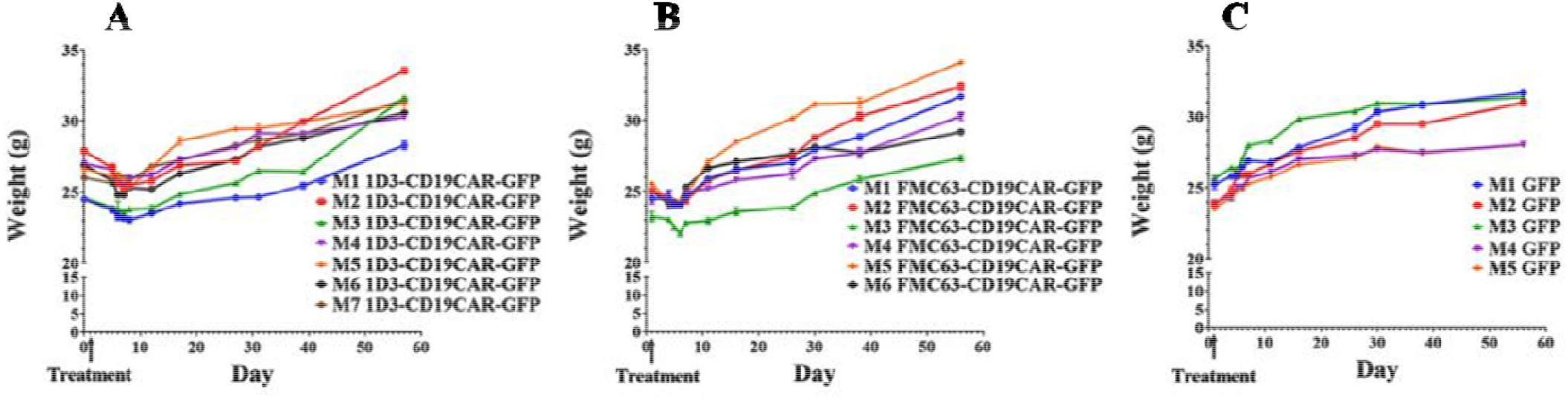
C57BL/6weights (grams) pre- and post treatment (n=8). Figure S6A: 1D3-CD19CAR-GFP lentivirus, Figure S6B. FMC63-CD19CAR-GFP lentivirus, and Figure S6C: GFP-only lentivirus or PBS. In **Figure S6A** and **S6B**, the mice that were treated with the 1D3-CD19CAR-GFP lentivirus or the FMC63-CD19CAR-GFP showed a 5.5 +/-2.97% (mean +/-SD) and 3.6 /-1.78% reduction in weight, respectively, within 5-7 days after receiving treatment. Once the mice weights recovered, they exhibited no other visible adverse effects.

**Figure S7.**
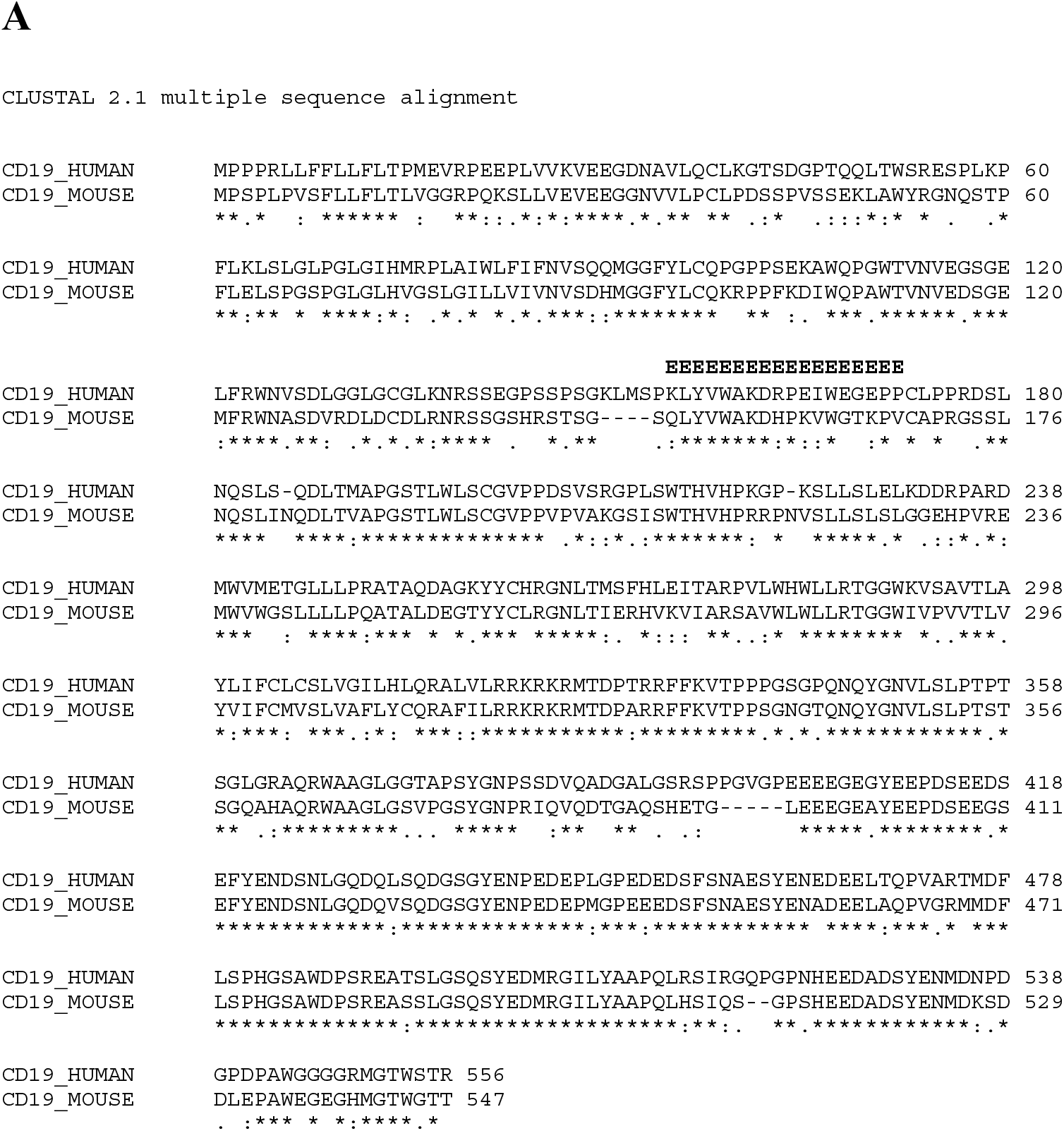
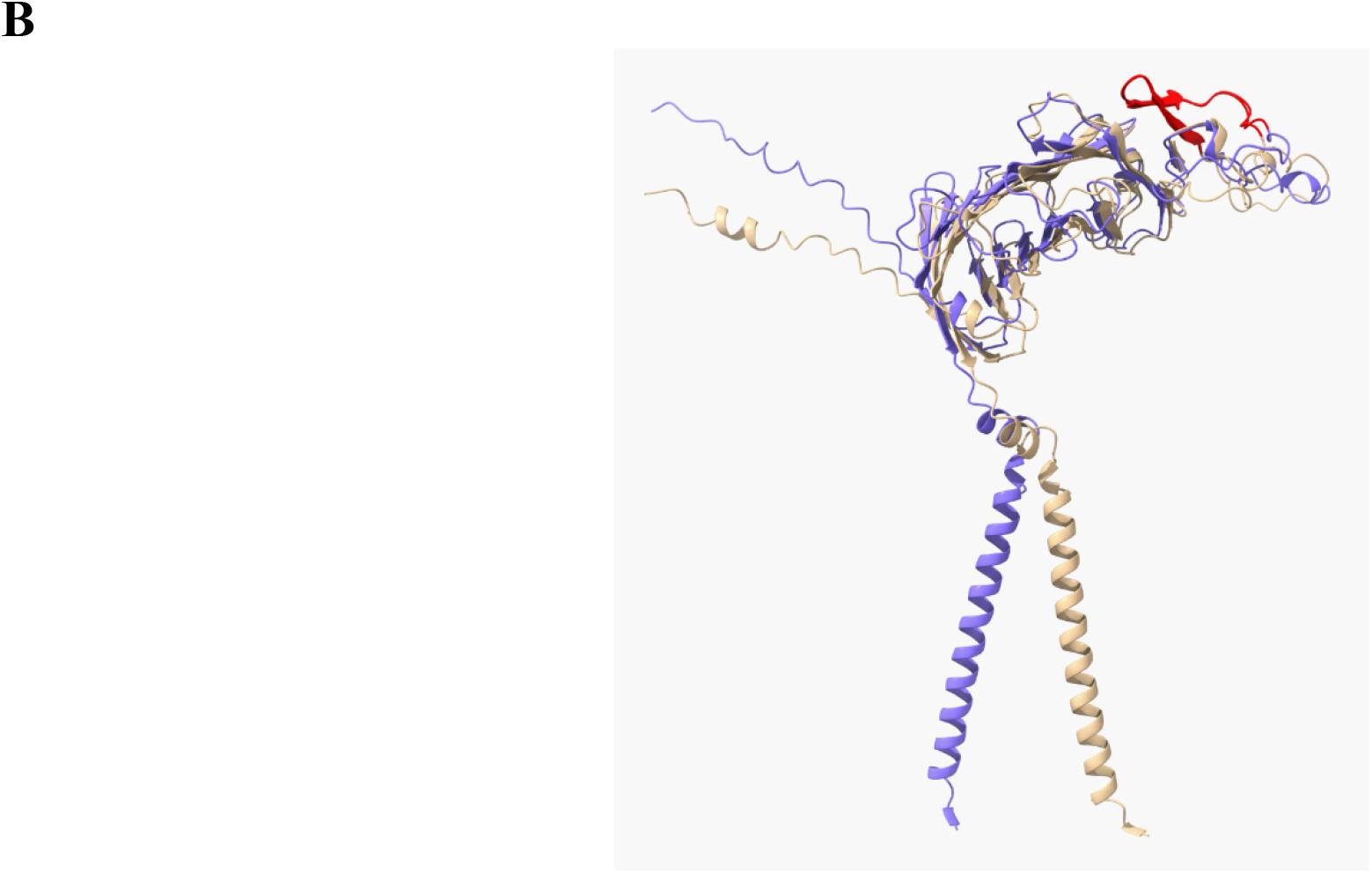
**(A)** Human and mouse CD19 multiple sequence alignment. CD19 protein sequences were obtained from Uniprot, and aligned using Clustal Omega with default settings. The region of the CD19 epitope (155-172 for CD19_HUMAN, 151-168 for CD19_MOUSE) is marked with ‘E’s above the sequence. **(B)** Human and mouse CD19 structure comparison. AlphaFold structure predictions for CD19 were obtained from Uniprot for human (tan) and mouse (blue). The epitope regions were aligned and colored red using ChimeraX. C-terminal polypeptides with poor AlphaFold prediction confidence scores were removed from the visualization (amino acids 331 to end for human, 329 to end for mouse). The structural similarity between the human and mouse CD19 proteins is high, especially in the area containing the epitope.

